# TRAF4 is crucial for the propagation of ST2^+^ memory Th2 cells in IL-33-mediated type 2 airway inflammation

**DOI:** 10.1101/2023.02.11.528093

**Authors:** Jianxin Xiao, Xing Chen, Weiwei Liu, Wen Qian, Katarzyna Bulek, Lingzi Hong, William Miller-Little, Xiaoxia Li, Caini Liu

**Author notes:** Corresponding authors: X.L., C.L.

## Abstract

Tumor necrosis factor receptor (TNF)-associated factor 4 (TRAF4) is an important regulator of type 2 responses in the airway; however, the underlying cellular and molecular mechanisms remain elusive. Herein, we generated T cell-specific TRAF4-deficient (CD4cre-*Traf4^fl/fl^*) mice and investigated the role of TRAF4 in interleukin (IL)-33 receptor (ST2, suppression of tumorigenicity 2)-expressing memory Th2 cells (ST2^+^ mTh2) in IL-33-mediated type 2 airway inflammation. We found that *in vitro* polarized TRAF4-deficient (CD4cre-*Traf4^fl/fl^*) ST2^+^ mTh2 cells exhibited decreased IL-33-induced proliferation as compared with TRAF4-sufficient (*Traf4^fl/fl^*) cells. Moreover, CD4cre-*Traf4^fl/fl^* mice showed less ST2^+^ mTh2 cell proliferation and eosinophilic infiltration in the lungs than *Traf4^fl/fl^* mice in the preclinical models of IL-33-mediated type 2 airway inflammation. Mechanistically, we discovered that TRAF4 was required for the activation of AKT/mTOR and ERK1/2 signaling pathways as well as the expression of transcription factor *Myc* and nutrient transporters (*Slc2a1, Slc7a1, and Slc7a5*), signature genes involved in T cell growth and proliferation, in ST2^+^ mTh2 cells stimulated by IL-33. Taken together, the current study reveals a previously unappreciated role of TRAF4 in ST2^+^ mTh2 cells in IL-33-mediated type 2 pulmonary inflammation, opening up avenues for the development of new therapeutic strategies.

**Graphical abstract:** 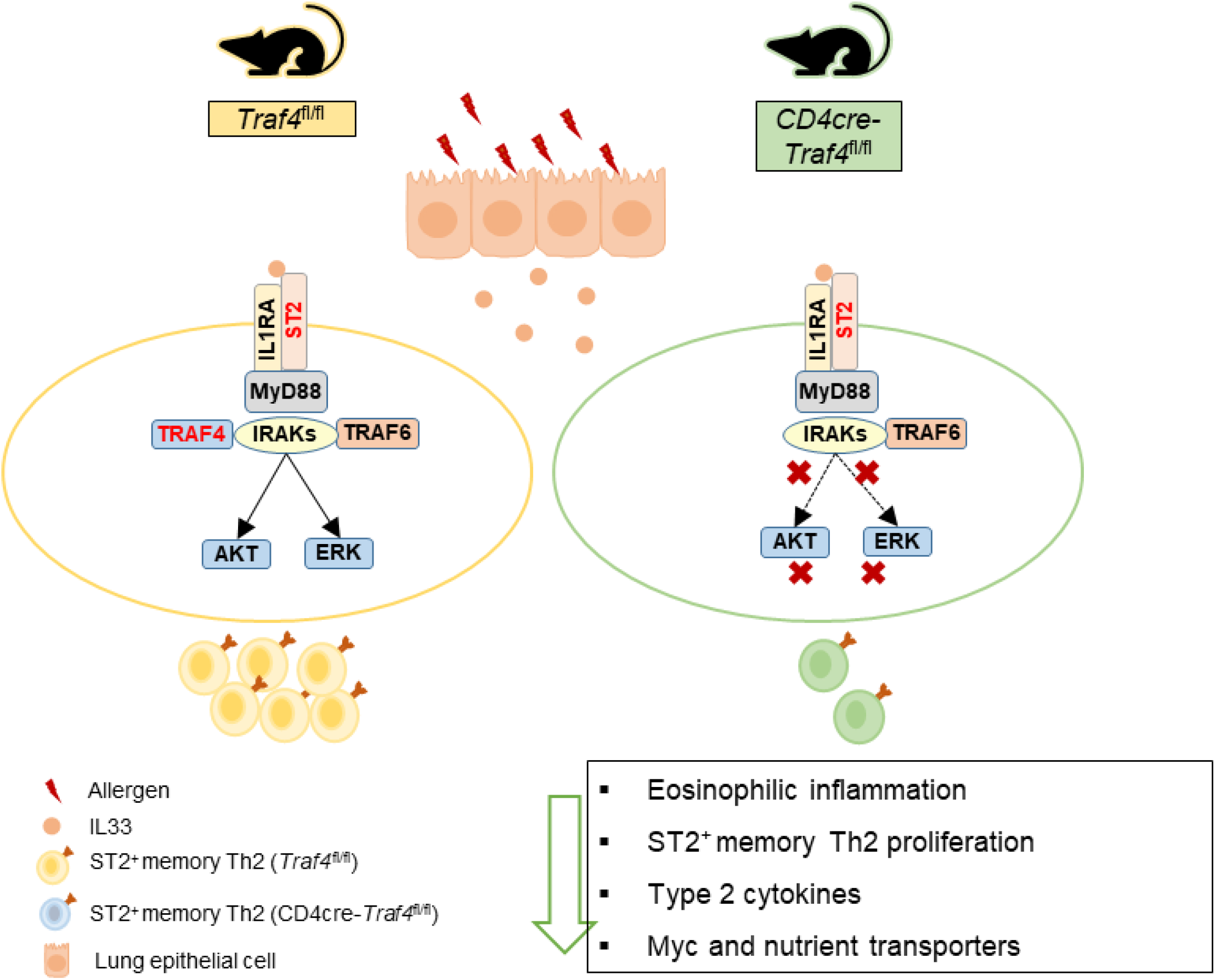

## Introduction

The alarmin cytokine interleukin (IL)-33, mainly produced by structural cells including endothelial, epithelial, and smooth muscle cells, is a well-known inducer and amplifier of type 2 immune responses in airway inflammation (1–5). IL-33 and its receptor subunit ST2 (suppression of tumorigenicity 2, encoded by *IL1RL1*) levels are elevated in asthmatic patients and mice and linked to the severity and steroid resistance of asthma (6–9). Furthermore, polymorphisms in *IL33* and *IL1RL1* have been associated with an increased risk of asthma in several genome-wide association studies involving a diverse range of geographical populations (1–5, 10). In addition, people carrying a loss-of-function mutation of *IL-33* have reduced blood eosinophil counts and are protected from asthma (11). Genetic ablation or antibody blockade of the IL-33/ST2 axis ameliorates airway inflammation in multiple preclinical models of allergic asthma (12–14). More interestingly, the results from two recent phase II clinical trials show that monoclonal antibodies targeting IL-33 (Itepekimab) or ST2 (Astegolimab) are beneficial to asthmatic patients by reducing blood eosinophils, improving lung function, or lowering exacerbation rate (15, 16).

IL-33 signals through a heterodimeric receptor complex consisting of ST2 and IL-1 receptor accessory protein (IL1RAcP). The activation of the IL-33 receptor complex leads to the recruitment of myeloid differentiation primary response protein 8 (MYD88), tumor necrosis factor (TNF) receptor-associated factor 6 (TRAF6), and IL-1 receptor-associated kinases (IRAKs), which results in activation of downstream pathways including mitogen-activated protein kinases (MAPKs), nuclear factor-κB (NF-κB) and AKT/mTOR, etc (17). IL33 targets multiple innate and adaptive cell types (e.g., group 2 innate lymphocytes (ILC2), mast cells, eosinophils, basophils, macrophages, dendritic cells, Th2, and Tregs) that are important players in type 2 immunity(17, 18).

IL-33-responsive ST2^+^ memory Th2 (mTh2) subsets play crucial roles in type 2 allergic airway inflammation. In humans, ST2-expressing mTh2 cells were confined or elevated in individuals with allergic airway diseases (19–21). In mice subjected to a house dust mite (HDM)-induced allergic asthma model, both circulating and resident ST2^+^ mTh2 (Trm-Th2) were shown to cooperatively promote the development of full-blown type 2 airway inflammation (22). ST2^+^ Trm-Th2 cells resided in a lymphoid-like structure called inducible bronchus-associated lymphoid tissue (iBALT) formed in the lung during chronic allergic inflammation and are maintained by IL-7- and IL-33-producing lymphatic endothelial cells (LECs)(23). Direct administration of IL-33 into mouse airways resulted in increased numbers of lung tissue-localized ST2^+^CD44^high^ mTh2 cells with enhanced production of IL-5 and IL-13 (9). Repetitive exposure of HDM to mice induced a significant increase in collagen deposition and increased numbers of ST2^+^ mTh2 cells producing amphiregulin (encoded by *Areg*, an important growth factor for tissue repair) in lung tissue (8). In an ovalbumin (OVA)-induced relapsing-remitting mouse model of allergic asthma, Th2 cytokine-producing ST2^+^CD44^+^CD69^+^ Trm-Th2 were found to mediate allergen-induced disease relapse and can maintain “allergic memory” in the lung for the lifetime of the mice(24).

The RING domain E3 ubiquitin ligase TRAF4 belongs to the TRAF family, which consists of seven members (TRAF1-7)(25–27). It acts either as a negative or positive regulator in multiple cytokine/growth factor signaling pathways (e.g. EGF, TGF beta, NOD2, IL-17 family, etc.), modulating cancer development and inflammatory diseases including asthma (25–28). TRAF4-deficient mice showed attenuated OVA- or IL-25 (IL-17E)-induced type 2 pulmonary inflammation, implicating that TRAF4 is an important regulator in these processes (29, 30). However, the cellular and molecular mechanisms of TRAF4 in type 2 immunity are still unclear.

In the current study, we generated T cell-specific TRAF4-deficient mice to investigate the intrinsic role of TRAF4 in ST2^+^ mTh2 cells in IL-33-mediated type 2 airway inflammation. The results from the present study indicate that TRAF4 is critical for the proliferation of ST2^+^ mTh2 induced by IL-33 in both *in vitro* and *in vivo* studies. Mechanistically, we demonstrated that TRAF4 is an essential signaling molecule for IL-33-mediated AKT/mTOR and ERK1/2 pathways, as well as for the expression of signature genes associated with T cell growth and proliferation in ST2^+^ mTh2 cells.

## Methods

### Mice

B6. OT-II TCR transgenic mice (Stock No: 004194) and CD4-Cre transgenic mice (Stock No: 022071) were purchased from the Jackson laboratories. *Rag2^-/-^/Il2rg^-/-^* mice (Model# 4111-F) were obtained from Taconic Biosciences. *Traf4* floxed (*Traf4^fl/fl^*) mice were generated by Cyagen Biosciences using gene-targeting technology. The experimental mice were female and age-matched (8-12 weeks). All mice were bred on C57BL/6 background and maintained under specific pathogen-free conditions. All animal experiments were performed according to The Cleveland Clinic’s IACUC.

### Reagents and cell culture

Mouse IL-2 (Cat# 575402), mouse IL-4 (Cat# 547302), mouse IL-33 (Cat# 580502), anti-mouse CD3ε (145-2C11), anti-mouse CD28 (37.51), anti-mouse IL-4 mAb (clone 11B11), anti-mouse IFN-γ mAb (XMG1.2) were purchased from Biolegend. Antibodies for TRAF4 (D1N3A), MyD88 (D80F5), phosphor (P)-mTOR (S2448, D9E), P-AKT (S473, D9E), P-AKT (T308, D25E6), P-JNK (T183/Y185, 81E11), P-p38 (T180/Y182, D3F9), P-ERK1/2 (T202/Y204, D13.14.4E), P-IκBα (S32, 14D4), and β-Actin (8H10D10) were obtained from Cell Signaling Technology. Mouse T1/ST2 antibody (DJ8) was from MD Bioproducts. AKT inhibitor VIII (CAS 612847-09-3) and LY3214996 (CAS 1951483-29-6) were purchased from Cayman. Primary mouse CD4^+^ T cells were cultured in complete TexMACS Medium (Miltenyi Biotec, #130-097-196) supplemented with 10% FBS, 50 μM β-mercaptoethanol, 100 U/ml penicillin, and 100 μg/ml streptomycin.

### In vitro polarization of ST2^+^ mTh2

Naïve CD4^+^ T cells (CD4^+^CD44^-^CD62L^+^) were isolated from mouse spleen by negative selection using the Naive CD4^+^ T Cell Isolation Kit (Miltenyi Biotec, 130-104-453). Purified naïve T cells were activated with plate-bound anti-CD3ε (5 μg/ml) and soluble anti-CD28 (2 μg/ml) and cultured in complete TexMACS Medium under Th2 polarizing conditions (50 ng/ml IL-4, 10 ng/ml IL-2, 10 μg/ml anti-IFN-γ) for 5 days. Then cells were washed with PBS and maintained in the medium with IL-2 and IL-7 (10 ng /ml, respectively) for 10 days to generate ST2^+^ mTh2 cells.

### IL-33-induced type 2 airway inflammation

Recombinant mouse IL-33 (1 μg/mouse) was intranasally (i.n.) administered in 40 μl sterile saline for 3 consecutive days. On day 10, BAL and lung tissue were harvested and analyzed.

### Alternaria-induced type 2 airway inflammation

*Alternaria alternata* (M1, Stallergenes Greer, 50 μg/mouse) dissolved in 40 μl sterile saline was administered (i.n.) on days 0, 6, and 9. BAL and lung were harvested and analyzed on day 12.

### ST2^+^ mTh2 adoptive transfer model

We bred TRAF4-sufficient *Traf4^fl/fl^* and TRAF4-deficient CD4-*Traf4^fl/fl^* mice onto OT-II TCR transgenic mice to obtain *Traf4^fl/fl^* OTII TCR and CD4-*Traf4^fl/fl^* OTII TCR transgenic mice. OVA_323–339_-specific ST2^+^ mTh2 cells were generated from naïve CD4^+^ T cells isolated from *Traf4^fl/fl^* OTII TCR and CD4-*Traf4^fl/fl^* OTII TCR as described above. Then these cells were transferred (i.v.) into 10-week female *Rag2^-/-^/Il2rg^-/-^* mice (10 × 10^6^ cells per mouse) and challenged (i.n.) with OVA_323–339_ (5 μg/ml) peptide with or without IL-33 (1 μg/mouse) for 3 consecutive days (day 0-2 after transfer). BAL cell counting and tissue collection were performed 3 days after the last antigen challenge.

### Real-time quantitative PCR (RT-PCR)

Total RNA was isolated using Quick-RNA Microprep Kit (ZYMO Research, Cat#R1050) as per the manufacturer’s instructions. First-strand cDNA was synthesized by ZymoScript RT PreMix Kit (ZYMO Research, Cat#R3012). All gene expression results were expressed as an arbitrary unit (AU) relative to the abundance of *Gapdh* mRNA. The primers used for the real-time-PCR are *Myc* (Forward: TCGCTGCTGTCCTCCGAGTCC; Backward: GGTTTGCCTCTTCTCCACAGAC), *Slc2a1* (Forward: GCTTCTCCAACTGGACCTCAAAC; Backward: ACGAGGAGCACCGTGAAGATGA), *Slc7a1* (Forward: CTCCTGGCTTACTCTTTGGTGG; Backward: GATCTAGCTCCTCGGTGGTTCT), *Slc7a5* (Forward: GGTCTCTGTTCACGTCCTCAAG; Backward: GAACACCAGTGATGGCACAGGT), and *Gapdh* (Forward: CATCACTGCCACCCAGAAGACTG; Backward: ATGCCAGTGAGCTTCCCGTTCAG).

### Periodic Acid Schiff (PAS) staining

Airway mucins were detected with PAS staining kit (Sigma-Aldrich, Cat# 395B-1KT). Sections of formalin-fixed, paraffin-embedded tissue were first deparaffinized, followed by incubation with Periodic Acid Solution (for 5 min) and Schiff’s Reagent (for 15 min), respectively. Then the slides were washed in running tap water for 3 min before counterstaining in Hematoxylin Solution (Sigma-Aldrich, Cat# 105174) for 10 s.

### Differential BAL cell counting

BAL cell counts were determined by cytospin slide preparation followed by Wright-Giemsa staining with Hema 3™ Manual Staining System and Stat Pack (Fisher Scientific, Cat# 22-122911)

### Flow cytometry and antibodies

Lung tissues were perfused by PBS and cut into small pieces before subjection to the digestion of Liberase TM (Millipore, Cat# 5401119001) plus DNase I (Millipore, Cat# 11284932001) for 45 min at 37 °C. The single-cell solution was obtained using a cell strainer (40 μm), followed by treatment with RBC lysis buffer (Biolegend, Cat# 420301). BAL cells and single lung cells were first gated by (FSC-A x SSC-A) to remove debris, followed by (FSC-A x FSC-H) to remove doublets, and then the dead cells were excluded using Zombie NIRTM Fixable Viability dye (Biolegend, Cat# 423105) (**Supplemental Figure 1, A and B**). The gated live cells were blocked by anti-mouse CD16/32 Ab (clone 2.4G, BD Biosciences, Cat# 553142) to avoid the non-specific binding to Fc receptors before surface or intracellular staining. Flow cytometry antibodies are from Biolegend [Alexa Fluor® 700 anti-mouse CD45 (30-F11), APC anti-mouse Ly-6G (1A8), PerCP/Cyanine5.5 anti-mouse CD44 (IM7), FITC anti-mouse Lineage Cocktail (anti-mouse CD3e, 145-2C11; anti-mouse Ly-6G/Ly-6C, RB6-8C5; anti-mouse CD11b, M1/70; anti-mouse CD45R/B220, RA3-6B2; anti-mouse TER-119/Erythroid cells, Ter-119), PE anti-mouse FOXP3 (150D), PE anti-mouse Ki-67 (16A8)], BD Biosciences [PE anti-mouse CD170 (Siglec-F, E50-2440), PE-CF594anti-mouse CD11b (M1/70), APC anti-mouse KLRG1 (MAFA)], Cell Signaling Technology [Myc (D84C12) mAb], and Thermo Fisher Scientific [PE-Cyanine7 anti-mouse IL13 (eBio13A); PE-eFluor® 610 anti-mouse IL13 (eBio13A), FITC anti-P-AKT1(S473) (AktS473-C7), PE Anti-P-4EBP (V3NTY24), APC Anti-P-S6 (cupk43k), PE-cy7 Anti-P-mTOR (MRRBY)]. Myc mAb (D84C12) was labeled by Alexa Fluor™ 647 Antibody Labeling Kit (Invitrogen, Cat# A20186). Surfaced staining was conducted using Cell Staining Buffer (Biolegend, Cat# 420201). Intracellular and nuclear staining was carried out with eBioscience™ Intracellular Fixation & Permeabilization Buffer (Cat# 88-8823-88) and eBioscience™ Foxp3 / Transcription Factor Staining Buffer Set (Cat# 00-5523-00) respectively. The sample analysis was performed using BD LSRFortessa Cell Analyzer.

### Cell proliferation assay

Cells were washed with PBS twice and labeled with CellTrace™ CFSE (ThermoFisher, Cat# C34554) (1 μM in PBS, 20 min, room temperature). Then the cells were incubated for 5 min with 5 volumes of complete TexMACS medium to remove any free dye before they were pelleted and suspended in fresh complete TexMACS medium for an additional 10 min and treated as indicated. The analysis was completed using BD LSRFortessa Cell Analyzer with FITC and Alexa Fluor 488 channel (488 nm excitation and a 530/30 nm bandpass emission filter).

### ELISA

Supernatants were collected from cell cultures and protein extracts of lung tissue were obtained using T-PER Tissue Protein Extraction Reagent (ThermoFisher, Cat# 78510). The levels of mouse IL-5 and IL-13 were measured with DuoSet ELISA kits (DY405 and DY413 respectively, R&D Systems). Mouse IL-33 was quantified by ELISA Kit (Cat# 88-7333-22) from Thermo Fisher Scientific.

### Co-immunoprecipitation (IP)

Cells were lysed on ice in Pierce™ IP Lysis Buffer (25 mM Tris-HCl pH 7.4, 150 mM NaCl, 1 mM EDTA, 1% NP-40, and 5% glycerol) (ThermoFisher, Cat# 87788) supplemented with cOmplete™ Protease Inhibitor Cocktail (MilliporeSigma, 11697498001), followed by centrifugation (10,000g, 10 min). The supernatant was immunoprecipitated using 2 μg/ml anti-mouse ST2 antibody and Pierce™ Protein G Magnetic Beads (ThermoFisher, Cat# 88848), followed by western blot analysis.

### Statistics

We used one-way or two-way ANOVA followed by Tukey’s multiple comparison test. P value (< 0.05) was indicated as significant. GraphPad Prism (V9.4) was used for all tests and calculations.

### Study approval

All animal studies were approved by The Cleveland Clinic’s IACUC.

## Results

### TRAF4 deficiency impairs IL-33-mediated proliferation of in vitro polarized ST2^+^ memory Th2

Previous studies show that TRAF4 is an important regulator of type 2 responses in murine allergic asthmatic models (29, 30). To explore the precise molecular and cellular mechanisms of TRAF4 in type 2 airway inflammation, we generated *Traf4^fl/fl^* mice with a floxed allele for *Traf4* by genetically targeting exon 2 of the *Traf4* (**Figure 1A**). To investigate the role of TRAF4 in T cells, we bred CD4cre transgenic mice onto *Traf4^fl/fl^* mice to generate T cell-specific TRAF4-deficient mice (CD4cre-*Traf4^fl/fl^*). Naive T cells (CD4^+^CD44^-^CD62L^+^) isolated from TRAF4-sufficient (*Traf4^fl/fl^*) mice and TRAF4-deficient (CD4cre-*Traf4^fl/fl^*) were cultured under Th2-polarizing condition (IL-2, IL-4, anti-IFN-γ Ab) in the presence of TCR stimulation (anti-CD3) and costimulation (anti-CD28) for 5 days to generate ST2^-^ effector Th2 cells (effTh2). Then effTh2 cells were removed from anti-CD3/anti-CD28 stimulation and cultured with IL-2 and IL-7 for additional 10 days to generate ST2^+^ mTh2 (**Figure 1B**). We found that both TRAF4-deficient (CD4cre-*Traf4^fl/fl^*) and control (*Traf4^fl/fl^*) cells differentiated into similar percentages of IL-13^+^ ST2^-^ effTh2 cells (57.2 vs 59.3) under IL-4/IL-2-driven Th2 polarizing condition and IL-13^+^ST2^+^ mTh2 cells (18.6 vs. 18.9) under IL-2/IL-7 culturing condition (**Figure 1C**). However, after IL-33 treatment of ST2^+^ mTh2 cells for another 3 days, we discovered that a higher percentage of ST2^+^ mTh2 cells were induced in *Traf4^fl/fl^* culture than in CD4cre-*Traf4^fl/fl^* culture (**Figure 1, C and D**). In addition, ST2^+^ cells in IL-33-treated CD4cre-*Traf4^fl/fl^* mTh2 culture displayed greater proliferating potential (as indicated by Ki-67 staining) as compared with ST2^+^ cells in IL-33-treated CD4cre-*Traf4^fl/fl^* mTh2 culture (**Figure 1, E and F**). Consistently, cell tracing experiments with CFSE also showed upon IL-33 stimulation, ST2^+^ mTh2 cells generated from *Traf4^fl/fl^* mice divided faster than ST2^+^ mTh2 from CD4cre-*Traf4^fl/fl^* mice (**Figure 1, G and H**). In contrast, the apoptosis rate of ST2^+^ cells was similar in TRAF4-sufficient (*Traf4^fl/fl^*) and TRAF4-deficient (CD4cre-*Traf4^fl/fl^*) mTh2 cells (**Figure 1, I and J**). IL-33-induced production of type 2 cytokines (IL-5 and IL-13) in culture supernatant were also greatly reduced in TRAF4-deficient (CD4cre-*Traf4^fl/fl^*) mTh2 cells as compared with those in control (*Traf4*^fl/fl CD4cre^) cells (**Figure 1K**). The above results indicate that TRAF4 is critical for IL-33-induced ST2^+^ mTh2 cell proliferation but dispensable for IL-2/IL-4-driven polarization of effTh2 and IL-2/IL-7-driven development of ST2-expressing mTh2.

**Figure 1.**
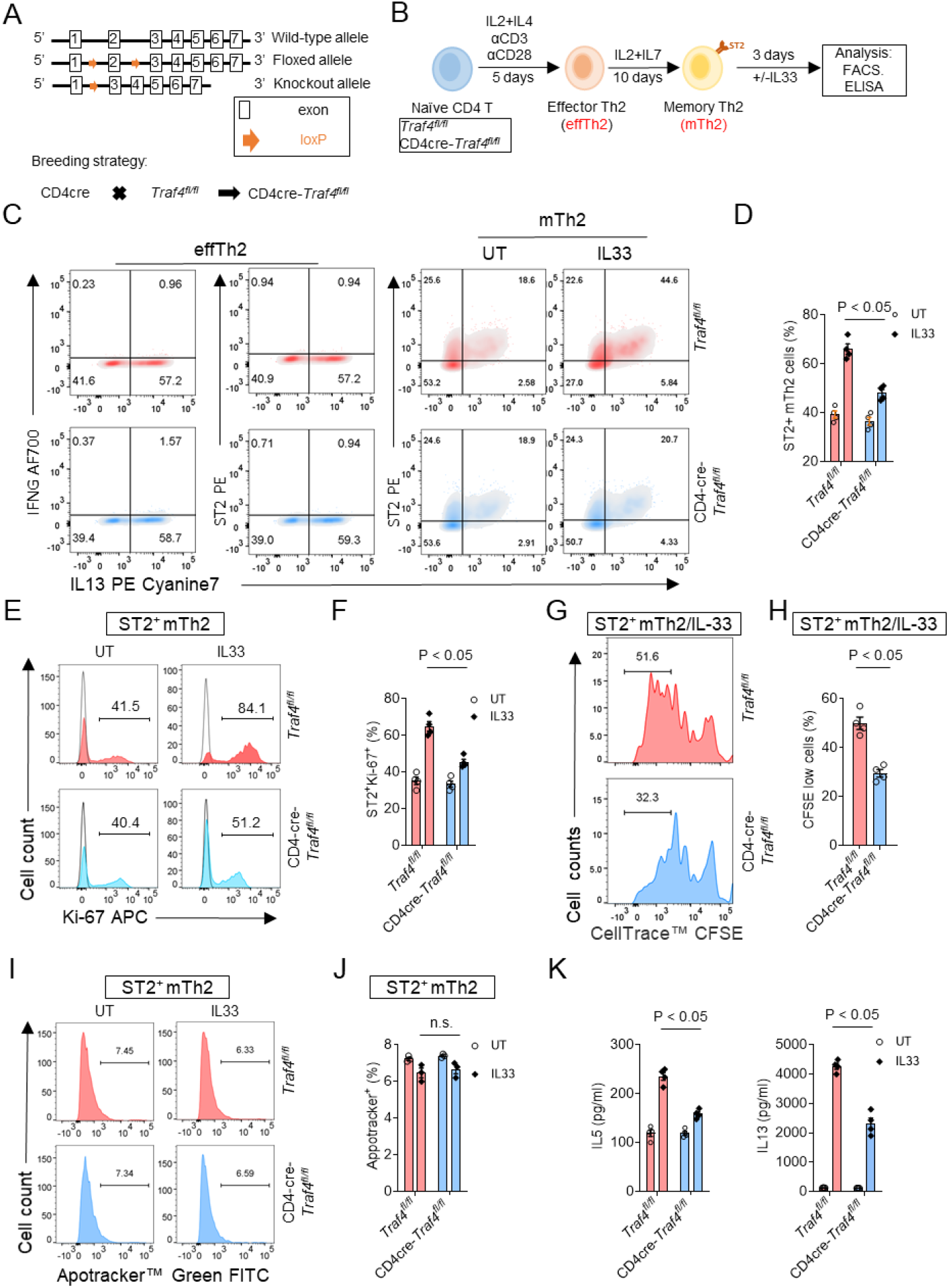
TRAF4 deficiency impaired IL-33-mediated proliferation of *in vitro* polarized ST2^+^ memory Th2. (A) Wild-type, floxed, and conditional knockout alleles of mouse *Traf4* are depicted schematically. (B) Naive CD4 T cells, isolated from T cell-specific TRAF4-deficient (CD4cre-*Traf4^fl/fl^*) and TRAF4-sufficient (*Traf4^fl/fl^*) mice, were differentiated under Th2 condition for 5 days to generate effector Th2 cells (effTh2); effTh2 were then cultured (with IL-2 and IL-7 only) for 10 days to obtain memory Th2 (mTh2). After that, mTh2 cells were then treated with sham (UT) or IL-33 for 3 days before indicated analysis (C-K). (C) Representative flow cytometry plots of effTh2 and sham- and IL-33-treated mTh2 cells. (D) Frequency of ST2^+^ mTh2 cells. (E) Histograms of Ki-67^+^ST2^+^ mTh2 cells. Filled histograms represent the cell population stained by the Ki-67 antibody, and the unfilled histograms represent the cell population stained by the isotype control antibody. (F) Frequency of Ki-67^+^ ST2^+^ mTh2 cells. (G) Histograms of IL-33-treated ST2^+^ mTh2 cells subjected to CFSE cell proliferation assay. (H) Frequency of CFSE low cells. (I) Histograms of apoptotic mTh2 cells (FITC^+^). (J) Frequency of apoptotic mTh2 cells. (K) IL-5 and IL-13 protein concentrations in cell medium were quantified by ELISA. Plotted data were shown as means ± SEM. Statistical analysis was performed with one-way ANOVA (Panel H) or two-way ANOVA (Panels D, F, G, K) followed by Turkey’s multiple comparison test. All data are representative of three independent experiments.

### T-cell-intrinsic TRAF4 is critical for the expansion of ST2^+^ Th2 memory cells in IL-33-induced airway inflammation

We next investigated the effect of T cell-specific TRAF4 deficiency on the proliferation of ST2^+^ CD4^+^ memory cells *in vivo*. We performed an IL-33 intranasal injection experiment on T cell-specific TRAF4-deficient (CD4cre-*Traf4^fl/fl^*) and control mice (*Traf4^fl/fl^*) (**Figure 2A**). As indicated by BAL counts [total leukocytes (CD45^+^) and eosinophils (CD45^+^CD11b^+^SiglecF^+^)] and lung histology, TRAF4-deficient (CD4cre-*Traf4^fl/fl^*) mice displayed reduced pulmonary inflammation as compared with TRAF4-sufficient (*Traf4^fl/fl^*) mice (**Figure 2, B and C**). The basal numbers of ST2^+^CD44^+^CD4^+^ (mTh2) cells in lung tissue (<1% of total lung CD4^+^ cells) were low in both CD4cre-*Traf4^fl/fl^* and *Traf4^fl/fl^* mice (**Figure 2E**). IL-33 induced greater expansion of ST2^+^CD44^+^CD4^+^ cells (8.5% of lung CD4^+^ cells) in *Traf4^fl/f^* mice than that (4.6% of lung CD4^+^ cells) in CD4cre-*Traf4^fl/fl^* mice (**Figure 2E**). Consistently, ST2^+^CD44^+^CD4^+^ cells from *Traf4^fl/fl^* mice had more proliferating potential (as indicated by the staining of cell proliferating marker Ki-67) than those from CD4cre-*Traf4^fl/fl^* mice (**Figure 2, E and F**). Since it has been known that IL-33 also induces the proliferation of ST2^+^Treg (FOXP3^+^) cells(31–34), we then further gated ST2^+^CD44^+^CD4^+^ cells into FOXP3^+^ (Treg) and FOXP3^-^ (mTh2) cells. Interestingly, both FOXP3^+^ and FOXP3^-^ cells were found to produce type 2 cytokine (as indicated by IL-13 intracellular staining), and IL-33-induced numbers of IL-13^+^ FOXP3^+^ and IL-13^+^FOXP3^-^ cells were higher in *Traf4^fl/fl^* mice than those in CD4cre-*Traf4^fl/fl^* mice (**Figure 2F**). We also examined IL-33-induced expansion of group 2 innate lymphocytes (ILC2, ST2^+^Lin^-^ KLRG1^+^), another major producer of type 2 cytokines in lung tissue exposed to IL-33 (35). In both IL-33-treated TRAF4-sufficient (*Traf4^fl/fl^*) and TRAF4-deficient (CD4cre-*Traf4^fl/fl^*) mice, we observed comparable numbers of IL-13^+^ILC2 (Lin-ST2^+^KLRG1^+^) and Ki67^+^ILC2 cells (**Figure 2, G and H**), implying that the reduced expansion of lung IL-13-producing ST2^+^CD44^+^CD4^+^ mTh2 cells in TRAF4-deficient (CD4cre-*Traf4^fl/fl^*) mice may be the cause of the attenuated eosinophilic airway inflammation. Consistently, IL-33-induced production of type 2 cytokines (IL-5 and IL-13) was also greatly reduced in lung tissue in TRAF4-deficient (CD4cre-*Traf4^fl/fl^*) mice as compared with control (*Traf4^fl/fl^*) mice (**Figure 2I**). Taken together, T-cell-specific TRAF4 deficiency impairs the expansion of IL-13-producing ST2^+^CD4^+^CD44^+^ mTh2 cells in the lung tissue and eosinophilic pulmonary inflammation in the IL-33 injection model.

**Figure 2.**
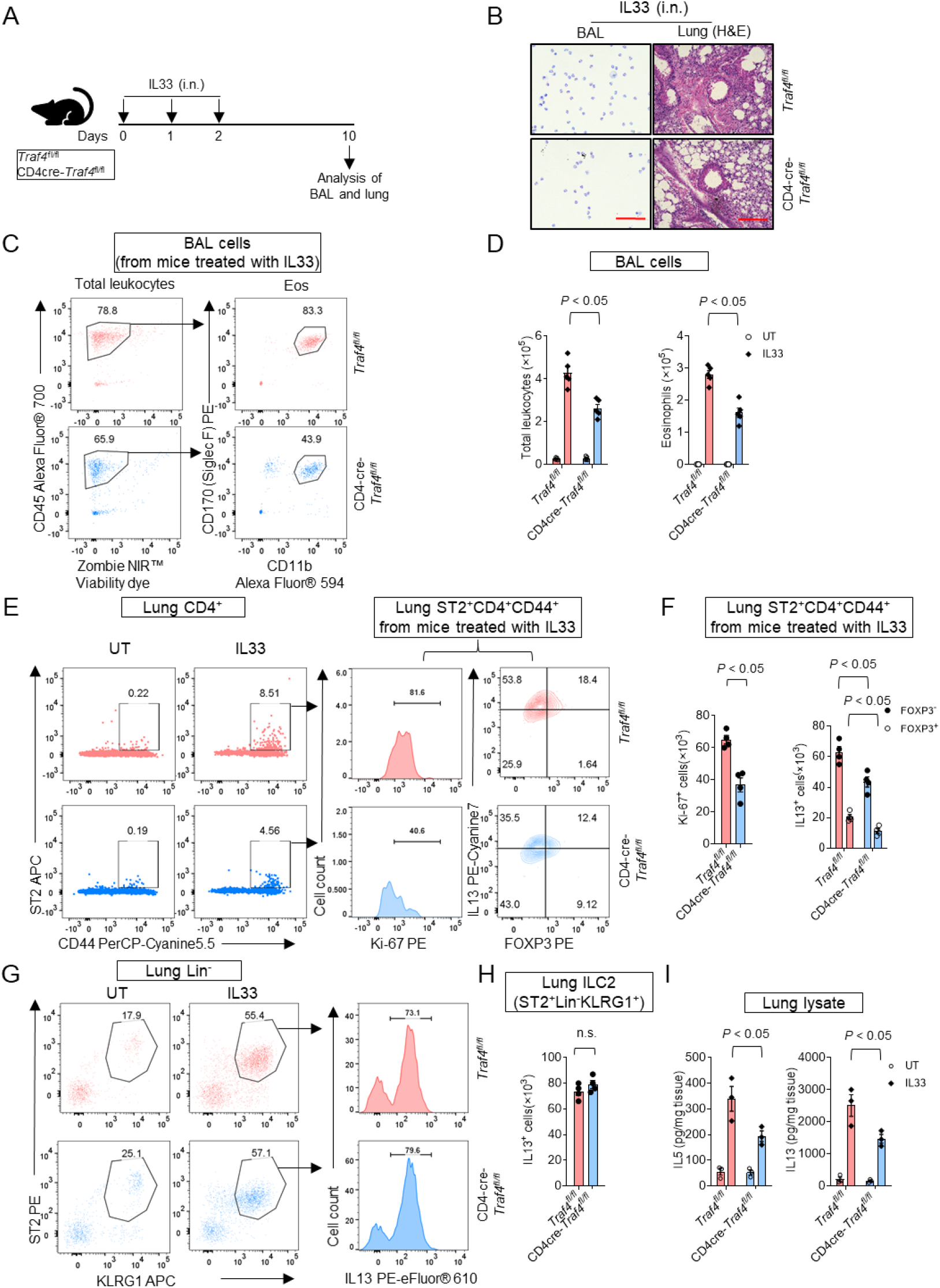
T cell-specific TRAF4 deficiency alleviated IL-33-induced type 2 airway inflammation. (A) Eight-week female T cell-specific TRAF4-deficient (CD4cre-*Traf4^fl/fl^*) and TRAF4-sufficient (*Traf4^fl/fl^*) were treated as indicated in the experimental protocol. (B-C) Giemsa staining of BAL cells and H&E staining of lung tissue section. Scale bars (red), 100 μm. (C) Flow cytometry plot of eosinophils (CD45^+^CD11b^+^SiglecF^+^) in the BAL. Eso, eosinophils. (D) Total leukocytes (CD45^+^) and eosinophils (CD45^+^CD11b^+^SiglecF^+^) in the BAL. (E) Flow cytometry analysis of lung CD4^+^ cells. (F) Absolute numbers of lung Ki-67^+^ST2^+^CD4^+^CD44^+^ and IL-13^+^ ST2^+^CD4^+^CD44^+^ cells. (G) Flow cytometry analysis of lung Lin^-^ (CD45^+^CD3^-^LY6G^-^LY6C^-^CD11b^-^ B220^-^TER-119^-^) cells. (H) Absolute numbers of lung Lin^-^ST2^+^KLRG1^+^ cells. (I) IL-5 and IL-13 protein concentrations in lung tissue were quantified by ELISA. Plotted data were shown as means ± SEM. Statistical analysis was performed with one-way ANOVA (Panels F and H) or two-way ANOVA (Panel I) followed by Turkey’s multiple comparison test. All data are representative of three independent experiments.

### T-cell-specific TRAF4 deficiency attenuates Alternaria-induced type 2 airway inflammation

We then sought to investigate the impact of T cell-specific TRAF4 deficiency on allergen-induced pulmonary inflammation. The fungal allergen (*Alternaria alternate*) is a rapid IL-33 inducer in the airways and is associated with asthma severity and exacerbations (36). We thus employed an *Alternaria*-driven type 2 asthma model in which IL-33 expression was significantly increased in lung tissue (**Figure 3, A and B**). *Alternaria* induced the expression of IL-33 at comparable levels in both *Traf4^fl/fl^* and CD4cre-*Traf4^fl/fl^* mice (**Figure 3B**). However, in comparison with TRAF4-sufficient control (*Traf4^fl/fl^*) mice, TRAF4-deficient CD4cre-*Traf4^fl/fl^* mice displayed greatly reduced eosinophilic inflammation [as indicated by total leukocytes (CD45^+^) and eosinophils (CD45^+^CD11b^+^SiglecF^+^) in the BAL] (**Figure 3C**) and lung histology (H&E and PAS staining) (**Figure 3D**)]. *Alternaria*-induced proliferation (as indicated by Ki-67 staining) of ST2^+^CD4^+^CD44^+^ cells as well as expansion of IL-13^+^ST2^+^CD4^+^CD44^+^ cells were greatly reduced in CD4cre-*Traf4^fl/fl^* mice as compared to control *Traf4^fl/fl^* mice (**Figure 3, E and F**). Following *Alternaria* challenge, the numbers of IL-13-producing mTh2 (Foxp3^-^ ST2^+^CD4^+^CD44^+^) and Treg (Foxp3^+^ ST2^+^CD4^+^CD44^+^) cells in lung tissue were also significantly lower in CD4cre-*Traf4^fl/fl^* mice than those in *Traf4^fl/fl^* mice (**Figure 3F**). In contrast, the number of IL-13^+^ILC2 (ST2^+^Lin^-^KLRG1^+^) was similar in both CD4cre-*Traf4^fl/fl^* and *Traf4^fl/fl^* mice (**Figure 3, G and H**). Additionally, we observed that TRAF4-deficient (CD4cre-*Traf4^fl/fl^*) lungs when exposed to *Alternaria* produced less amounts of type 2 cytokines (IL-5 and IL-13) than control (*Traf4^fl/fl^*) lungs, which was in line with the changes in pulmonary pathology (**Figure 3I**). It should be noted that lung IL-13^+^ST2^+^CD4^+^CD44^+^ mTh2 cells outnumbered IL-13^+^ ILC2 cells by more than five times in TRAF4-sufficient *Traf4^fl/fl^* mice subjected to the *Alternaria* model (**Figure 3, F and H**), indicating that IL-13^+^ST2^+^CD4^+^CD44^+^ mTh2 cells contribute more to *Alternaria*-induced airway inflammation than IL-13^+^ ILC2 cells. This is in contrast with the almost equivalent numbers of IL-13^+^ST2^+^CD4^+^CD44^+^ mTh2 and IL-13^+^ ILC2 cells in the lung tissue from the *Traf4^fl/fl^* mice subjected to IL-33 injection (**Figure 2, F and H**). As a result, the mice on *Alternaria* model exhibited a more dramatic phenotype caused by T cell-specific TRAF4 deficiency than the mice on IL-33 injection model (**Figure 2, B-D; Figure 3, C and D**). The above results indicate that T-cell-specific TRAF4 is crucial for allergen-induced eosinophilic airway inflammation and expansion of IL-13^+^ST2^+^CD4^+^CD44^+^ mTh2 cells in the lung.

**Figure 3.**
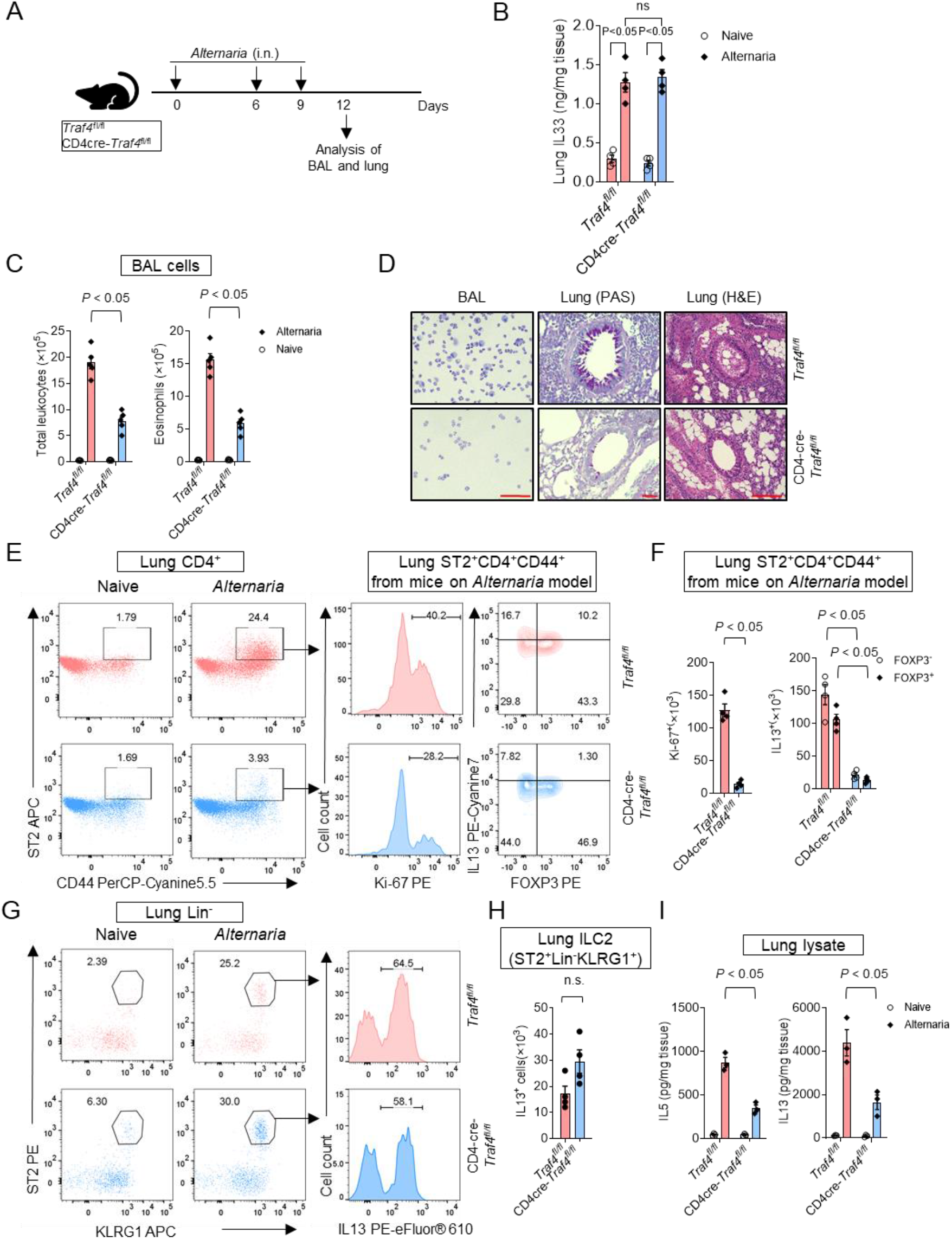
T cell-specific TRAF4 deficiency mitigated *Alternaria*-induced type 2 airway inflammation. (A) Ten-week female T cell-specific TRAF4-deficient (CD4cre-*Traf4^fl/fl^*) and TRAF4-sufficient (*Traf4^fl/fl^*) mice were treated as indicated experimental protocol. (B) Lung IL-33 protein concentrations were measured by ELISA. (C) Total leukocytes (CD45^+^) and eosinophils (CD45^+^CD11b^+^SiglecF^+^) in the BAL. (D) Giemsa staining of BAL cells and H&E staining of lung tissue section. Scale bars (red), 100 μm. (E) Flow cytometry analysis of lung CD4^+^ cells. (F) Absolute numbers of lung Ki-67^+^ST2^+^CD4^+^CD44^+^ and IL-13^+^ST2^+^CD4^+^CD44^+^ cells. (G) Flow cytometry analysis of lung Lin^-^ cells. (H) Absolute numbers of lung Lin^-^ST2^+^KLRG1^+^ILC2 cells. (I) Lung IL-5 and IL-13 protein levels were measured by ELISA. Plotted data were shown as means ± SEM. Statistical analysis was performed with one-way ANOVA (Panels F and H) or two-way ANOVA (Panel I) followed by Turkey’s multiple comparison test. All data are representative of three independent experiments.

### The intrinsic TRAF4 is crucial for IL-33-induced expansion of OVA_323-339_-specific ST2^+^ mTh2 cells and the development of eosinophilic airway inflammation in the adoptive transfer mice

Since IL-33- and *Alternaria*-induced type 2 inflammation involves induction of both ST2^+^ mTh2 and ST2^+^ILC2 lymphocytes (**Figure 2, F and H; Figure 3, F and H**), we sought an *in vivo* model in which the ST2^+^ mTh2 cells are the solely lymphocyte population driving type 2 airway inflammation. We performed the adoptive transfer experiment with *Rag2^-/-^IL2rg^-/-^* (R2G2) mice (genetically lacking ILCs, T cells, B cells, and NK cells) as recipients (**Figure 4A**). In this experimental model, R2G2 mice were subjected to adoptive transfer of OVA_323-339_-specific TRAF4-sufficient (*Traf4^fl/fl^*) or TRAF4-deficient (CD4-*Traf4^fl/fl^*) ST2^+^ mTh2 cells so that the impact of ST2^+^ mTh2 (not ST2^+^ ILC2 and ST2^+^ Tregs) was manifest. After transfer, the R2G2 recipient mice were challenged (i.n.) with the low dose (5 μg) of OVA_323-339_ peptide along with sham or IL-33 treatment. OVA_323-339_ peptide induced similar low-grade eosinophilic airway inflammation [as indicated by the numbers of eosinophils (CD45^+^CD11b^+^SiglecF^+^) in the BAL) of both TRAF4-sufficient (*Traf4^fl/fl^*) and TRAF4-deficient (CD4cre-*Traf4^fl/fl^*) mice (**Figure 4, B and C**). However, after being co-treated with IL-33, recipient mice transferred with TRAF4-sufficient (*Traf4^fl/f^*) cells developed significantly more eosinophilic inflammation in the lung than the mice transferred with TRAF4-deficient (CD4cre-*Traf4^fl/f^*) cells (**Figure 4, B and C**). Likewise, the numbers of proliferating ST2^+^CD4^+^ cells (Ki-67^+^) and cytokine-producing IL-5^+^IL-13^+^ST2^+^CD4^+^ cells in IL-33/OVA_323-339_-treated mice transferred with TRAF4-sufficient (*Traf4^fl/fl^*) cells were more than those in IL-33/OVA_323-339_-treated mice transferred with TRAF4-deficient (CD4cre-*Traf4^fl/fl^*) cells (**Figure 4, D and F**). The data presented above suggest that the intrinsic TRAF4 deficiency impairs the IL-33-induced expansion of antigen-specific ST2^+^ mTh2 as well as the development of eosinophilic inflammation in the absence of ST2^+^ ILC2 and other lymphocytes in the lungs of adoptive transfer mice.

**Figure 4.**
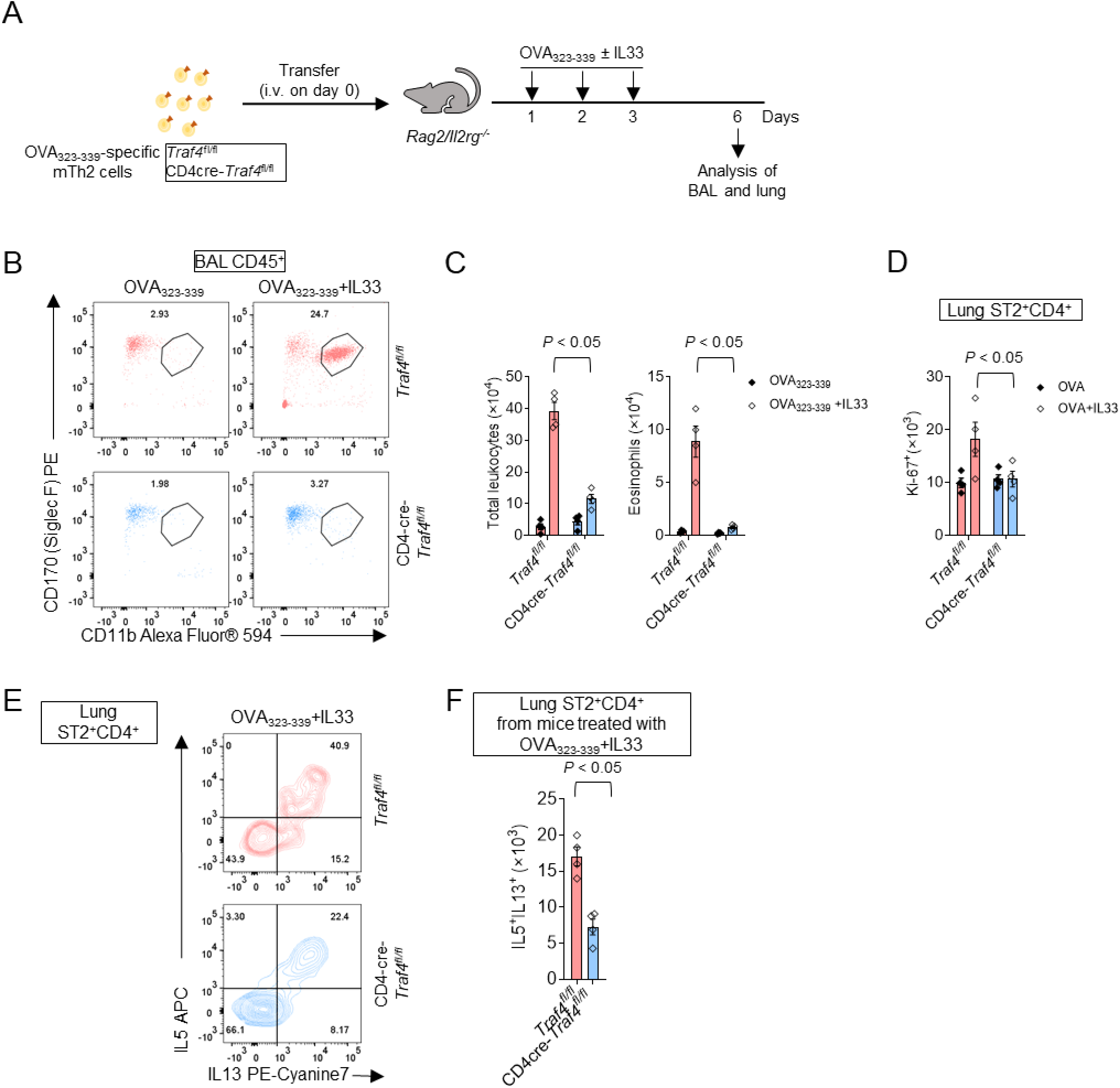
The intrinsic TRAF4 is crucial for IL-33-induced expansion of OVA323-339-specific ST2^+^ mTh2 and the development of eosinophilic airway inflammation in the adoptive transfer mice. (A) Ten-week female *Rag2/Il2rg^-/-^* mice were adoptively transferred with T cell-specific TRAF4-deficient (CD4cre-*Traf4^fl/fl^*) and TRAF4-sufficient (*Traf4^fl/fl^*) mTh2 cells and challenged with OVA peptide with or without IL-33 as indicated in the experimental protocol. (B) Flow cytometry analysis of BAL eosinophils (CD45^+^CD11b^+^SiglecF^+^). (C) Total leukocytes (CD45^+^) and eosinophils (CD45^+^CD11b^+^SiglecF^+^) in the BAL. (D) Flow cytometry analysis of lung Ki-67^+^ST2^+^CD4^+^ cells. (E) The absolute number of lung IL-13^+^IL-5^+^ ST2^+^CD4^+^ cells. Plotted data were shown as means ± SEM. Statistical analysis was performed with one-way ANOVA (Panel F) or two-way ANOVA (Panels C and D) followed by Turkey’s multiple comparison test. All data are representative of three independent experiments.

### TRAF4 is required for IL-33-mediated PI3K/AKT and ERK1/2 pathways as well as signature genes involved in signature genes involved in T cell growth and proliferation in ST2^+^ mTh2 cells

We next investigated the molecular mechanism of TRAF4 in IL-33-induced ST2^+^ mTh2 proliferation. We performed a co-immunoprecipitation (IP) experiment with an antibody targeting endogenous IL-33 receptor (ST2) using *in vitro* polarized ST2^+^ mTh2 cells. As shown in **Figure 5A**, upon IL-33 stimulation, TRAF4 was recruited to ST2 along with known signaling molecular MYD88, indicating that TRAF4 forms a proximal receptor complex with it. It is known that IL-33 induces multiple pathways (e.g. NF-κB, MAPKs, PI3K/AKT/mTOR) in different types of cells (17). We then examined the activation of these signaling pathways by IL-33 in ST2^+^ mTh2 cells cultured from TRAF4-sufficient (*Traf4^fl/fl^*) and TRAF4-deficient (CD4-*Traf4^fl/fl^*) mice (**Figure 5, B and C**). We found that IL-33-induced activation of AKT/mTOR and ERK1/2 (as shown by the phosphorylation of the indicated signaling molecules) was markedly reduced in TRAF4-deficient (CD4-*Traf4^fl/fl^*) cells as compared with that in TRAF4-sufficient (*Traf4^fl/fl^*) cells. TRAF4 deficiency also diminished IL-33-induced phosphorylation of ribosomal protein S6 (P-S6) and eukaryotic initiation factor 4E binding protein 1 (P-4EBP1), key downstream effectors of mTOR for protein synthesis (**Supplemental Figure 2, A and B**). In contrast, TRAF4 deficiency did not affect the activation of JNK and p38 MAPKs (as shown by the phosphorylation of the indicated kinases) by IL-33, while the activation of NF-κB (as indicated by P-IκBα) was negatively regulated by TRAF4 deficiency. We then assessed the effect of AKT inhibitor VIII (an allosteric inhibitor of AKT1 and AKT2) (37, 38) and LY3214996 (a selective ERK1/2 inhibitor) (39) on IL-33-induced ST2^+^ mTh2 proliferation by a cell tracing experiment with CFSE. As shown in **Figure 5, D and E**, IL-33-induced mTh2 proliferation was greatly reduced by AKT inhibitor VIII and attenuated by LY3214996. These findings suggest that TRAF4 regulates IL-33-induced mTh2 proliferation via the AKT/mTOR and ERK1/2 pathways. We also compared the IL-33-induced expression of the signature genes involved in T cell growth and proliferation between TRAF4-sufficient (*Traf4^fl/fl^*) and TRAF4-deficient (CD4-*Traf4^fl/fl^*) ST2^+^ mTh2 cells (**Figure 5F**). We found that the transcripts of *Myc* (the master transcription factor for TCR-induced T cell activation), *Scl2a1* (glucose transporter protein type 1, GLUT1), *Scl7a1* (cationic amino acid transporter 1, CAT1), and *Scl7a5* (L-type amino acid transporter 1, LAT1) were noticeably upregulated by IL-33 stimulation. However, TRAF4 deficiency did not affect the mRNA expression of *Myc* and *Scl2a1* but impaired transcription of *Scl7a1* and *Scl7a5* in ST2^+^ mTh2 after IL-33 stimulation for 24 h. Interestingly, TRAF4 deficiency substantially reduced the intracellular protein level of *Myc* and the surface protein expression of *Scl2a1* upregulated by IL-33 (**Figure 5, G and H**). The above results suggest that a subset of IL-33-induced genes involved in cell growth and proliferation are TRAF4-dependent in ST2^+^ mTh2 cells.

**Figure 5.**
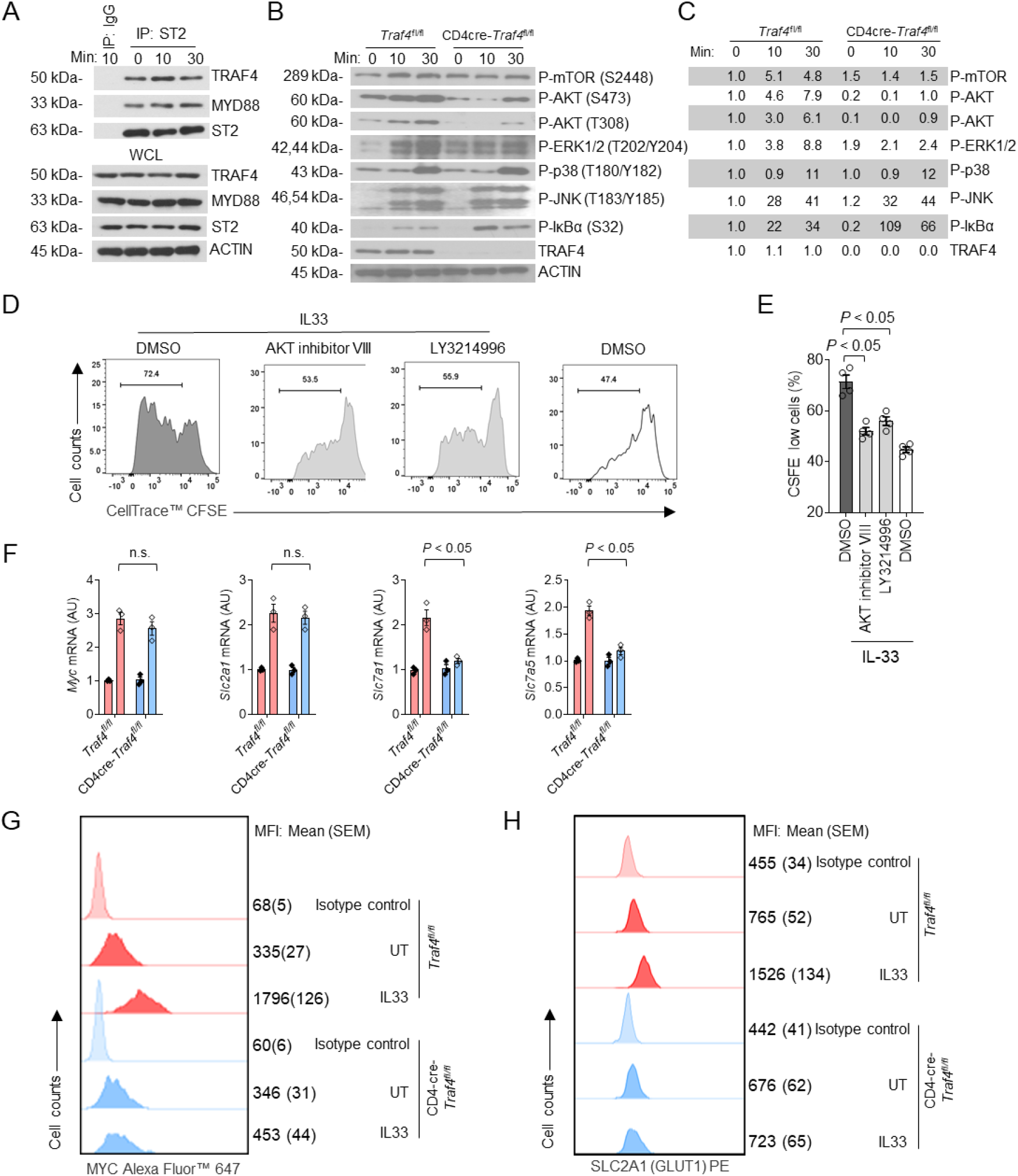
TRAF4 is required for IL-33-mediated AKT/mTOR and ERK1/2 pathways as well as signature genes involved in T cell growth and proliferation in ST2^+^ mTh2 cells. (A) Cell lysates of *in vitro* polarized mTh2 treated with or without IL-33 were subjected to the co-immunoprecipitation assay with anti-ST2 antibody, followed by western blot analysis with the indicated antibodies. (B) mTh2 cells (starved in cytokine free medium for 24 h) were treated with IL-33 at different time points, and cell lysates were analyzed by western blot with indicated antibodies. (C) The density of each protein band was quantified by Image J (V. 1.51) and normalized to that of actin. Then the fold induction was calculated related to the value of *Traf4^fl/fl^* sample at time 0 min (which was set to 1) for each protein. (D) Histograms of ST2^+^ mTh2 cells subjected to CFSE cell proliferation assay. (E) Frequency of CFSE low cells. (F) Real-time PCR analysis of the mRNA abundance in mTh2 after 24 h treatment. (G-H) Representative histogram blots showing the intracellular expression of MYC and the surface expression of GLUT1 on mTh2 cells treated with sham or IL-33 for 24 h. MFI, Mean fluorescence intensity. Plotted data were shown as means ± SEM. Statistical analysis was performed with one-way ANOVA (Panel E) or two-way ANOVA (Panel F) followed by Turkey’s multiple comparison test. All data are representative of two independent experiments.

## Discussion

Asthma is a prevalent immunologic disorder, around half of which is driven by type 2 inflammation and characterized by increased eosinophilic infiltration in the lung. Although the majority of type 2 asthmatics are responsive to steroid-based therapy, a subgroup of patients, especially those with persistent eosinophilic inflammation, are refractory to steroid treatment and require high-dose steroids to control the symptoms(40). Increased IL-33 expression is associated with severe and refractory asthma (6, 7). IL-33 administration to the mouse airway induces the expansion and cytokine production of both ST2^+^ mTh2 cells and ST2^+^ ILC2, contributing to antigen-dependent and -independent type 2 airway inflammation (41). Steroid-resistant (SR) subsets of ST2^+^ mTh2 and ST2^+^ ILC2 have been reported in allergic human patients or mice on experimental models of eosinophilic airway diseases (9, 42-44). Targeting IL-33-induced ST2^+^ mTh2 and ST2^+^ ILC2, the major producers of type 2 cytokines in the airway, may be effective for SR eosinophilic airway inflammation. Interestingly, two recent phase II clinical trials have indicated that mAbs blocking either IL-33 or its receptor ST2 were beneficial for certain subsets of asthmatic patients (15, 16). The current study identified TRAF4 as a crucial regulator of the proliferation of ST2^+^ mTh2 induced by IL-33 in *in vitro* cell culture studies as well as in *in vivo* murine models of type 2 airway inflammation, which may open up avenues for the development of new therapeutic strategies for SR eosinophilic asthma.

TRAF4 has been reported as a positive regulator for cell growth/proliferation pathways (AKT, ERK1/2, and ERK5) induced by EGF, TGFβ, and IL-17A signaling, contributing to the proliferation and migration of cancer cells (45–48). AKT/mTOR and ERK1/2 pathways are also critical for CD4 T cell proliferation in response to TCR activation and IL-2 (49). In the current study, we found that TRAF4, along with MYD88, is recruited to the IL-33 receptor (ST2) upon IL-33 stimulation, forming a proximal signaling complex in ST2^+^ mTh2 cells. Furthermore, TRAF4 deficiency impaired IL-33-induced activation of AKT/mTOR and ERK1/2 pathways as well as the proliferation of ST2^+^ mTh2 cells. TRAF4 is known to bind phosphatidylinositol phosphate (PIP) lipids (e.g. PIP2 and PIP3) on cell membranes (50), which may facilitate PI3K kinase recruitment and the activation of downstream AKT/mTOR pathway. Conversely, we also discovered that TRAF4 negatively regulates the phosphorylation of IκB, an important upstream event promoting NF-κB translocation into the nucleus and activation. Activated IL-33 receptor recruits IRAK kinases, which bind and activate TRAF6 (a well-known upstream signaling molecule for the NF-κB pathway)through TRAF-binding sites (51). There likely exists a competition between TRAF4 and TRAF6 for TRAF-binding sites on IRAKs in the IL-33 pathway. A similar mechanism has been depicted in the IL-17A pathway in which TRAF4 and TRAF6 compete for the TRAF-binding motifs on Act1 (the essential signaling adaptor for IL-17A receptor) (52).

Prior research has highlighted the innate (TCR-independent) immune functions of ST2^+^ mTh2 cells. *In vitro* polarized ST2^+^ mTh2 can produce IL-5 and IL-13 upon IL-33 stimulation alone or with a STAT5 activator (e.g. IL-2, IL-7, or TLSP) (53). *In vivo* studies also revealed that ST2^+^ mTh2 cells in lung tissue produced type 2 cytokines in response to IL-33 or IL-33-inducing allergens in a TCR-independent manner(54). In the current study, we also observed that IL-33 was able to induce the production of IL-5 and IL-13 in our *in vitro* polarized ST2^+^ mTh2 cells in the presence of IL-2 and IL-7. Moreover, the administration of IL-33 and IL-33-inducing allergen (e.g. *Alternaria*) to the airway was able to induce the expansion of IL-13^+^ST2^+^CD4^+^ mTh2 cells in lung tissue. Therefore, our studies provide additional evidence to support the innate functions of ST2^+^ mTh2 in antigen-independent type 2 airway inflammation.

ST2^+^ Tregs (characterized by FOXP3 expression) is an important subset of CD4 T cells that are notably found at barrier sites (e.g. gut, lung). ST2^+^ Tregs exert either suppressive or proinflammatory functions in different experimental contexts. In respiratory infection models, IL-33 maintained FOXP3 expression and suppressive function of Tregs, which inclined to acquire Th17 phenotype (upregulation of RORC and IL-17 expression) in the absence of ST2(34). In a mouse model of T-cell-induced colitis, ST2 expression by Tregs was critical to prevent the onset of disease in the gut. Conversely, the additional studies showed that ST2^+^ Tregs exhibited Th2-biased characteristics, producing type 2 cytokines (IL-5 and IL-13) and even losing their ability to suppress effector T cells when stimulated *in vitro* with IL-33(33, 55). Airway administration of IL-33 to mice impaired established immunologic tolerance to the ovalbumin antigen, which was accompanied by an increased number of ST2^+^ Tregs expressing IL-5 and IL-13 (33). Interestingly, in an HDM-induced type 2 pulmonary inflammation model, ST2^+^ Tregs activated by IL-33 were able to suppress IL-17-producing gamma/delta T cells but not adaptive Th2 immune response(56). The findings from the current study discovered that airway administration of IL-33 or IL-33-inducing allergen (*Alternaria*) promoted the expansion of IL-13-expressing FOXP3^-^ (mTh2) and FOXP3^+^ (Treg) ST2^+^ CD4 cells in the lung, suggesting that both mTh2 and Tregs may contribute to airway inflammation as the producers of type 2 cytokines. Additionally, we found that TRAF4 deficiency impaired IL-33-induced propagation of both FOXP3^-^ mTh2 and FOXP3^+^ Tregs. However, the precise role of TRAF4 in Tregs needs to be further investigated using FOXP3-specific TRAF4-deficient mice in type 2 airway inflammation.

The transcription factor MYC is considerably increased after TCR ligation, and this promotes the transcription of vital enzymes and transporters involved in a variety of metabolic pathways during the activation and growth of CD4 T cells. (57, 58). MYC deficiency drastically decreases T cell growth and proliferative capacity (57–59). Multiple mechanisms, including transcriptional, post-transcriptional, and post-translational controls, are involved in the regulation of MYC expression (60–67). In particular, MYC protein level is regulated by phosphorylation/dephosphorylation-mediated stabilization and degradation events (64, 65, 67). ERK activity stabilizes MYC by phosphorylation at serine 62, whereas glycogen synthase kinase 3β destabilizes it by phosphorylation at threonine 58, which can be blocked by the PI3K/AKT pathway(59, 64, 68). In the present study, we observed that TRAF4 deficiency had little impact on MYC transcription but greatly reduced MYC protein level in IL-33-stimulated ST2^+^ mTh2, suggesting that TRAF4 may affect the posttranslational control of MYC protein through ERK1/2 and PI3K/AKT pathways.

MYC-mediated induction of amino acid transporters is pivotal for the bioenergetic and biosynthetic pathways in TCR-activated T cells (57). In the present study, IL-33-induced transcription of two MYC-dependent amino acid transporters (*Scl7a1* and *Slc7a5*) was impaired in TRAF4-deficient ST2^+^ mTh2 cells. *Slc7a1* encodes cationic amino acid transporters 1 (CAT1), which is a key transporter for L-arginine and L-ornithine. Arginases convert L-arginine to L-ornithine, which is an important precursor for the synthesis of polyamines (spermidine, spermine, and putrescine). Polyamine metabolites have been associated with cell growth and proliferation (69). Arginase 1 (ARG1) deficiency in ILC2 leads to reduced polyamine synthesis and impaired cell proliferation (70). Likewise, the reduced surface expression of *Scl7a1* in TRAF4-deficient ST2^+^ mTh2 cells may impair the uptake of arginine, leading to reduced polyamine synthesis and cell proliferation. *Scl7a5* encodes the light chain unit of the large neutral amino acid (LNAA) transporter 1 (LAT1), which is the main transporter for LNAAs like leucine and methionine. Leucine is an important nutrient signal for mTORC1 activation (71), and methionine (the starting amino acid for each protein) availability sets the limit for the overall efficiency of protein synthesis. *Slc7a5^−/−^* T cells have defective mTOR activation and proliferation upon TCR ligation, which recapitulates the phenotype of *Myc*^-/-^ T cells (57, 72). The current study indicates that IL-33, similar to TCR ligation, induces MYC-dependent expression of amino acid transporters to meet the increased metabolic needs for the growth and proliferation of ST2^+^ mTh2 cells. It will be interesting to map the complete proteomes of ST2^+^ mTh2 reprogrammed by IL-33 stimulation and understand the effect of TRAF4 deficiency on the overall amino acid transport.

The present study found that surface expression of glucose transporter GLUT1 was greatly attenuated by TRAF4 deficiency in IL-33-treated ST2^+^ mTh2 cells although the transcript abundance of GLUt1 was not affected. Despite MYC being previously reported to be critical for mRNA expression of GLUT1 of T cells upon TCR activation (58), more recent studies indicate that GLUT1 transcription is MYC-independent (57). Interestingly, intracellular trafficking of GLUT1 has been shown to depend on PI3K/AKT pathway in different cell types (73–75). Therefore, the impaired AKT pathway in TRAF4-deficient ST2^+^ mTh2 may impede the trafficking of GLUT1, resulting in the tapered surface expression of GLUT1. Future research should interrogate the specifics of how TRAF4 deficiency impacts IL-33-induced glucose metabolic pathways in ST2^+^ mTh2 cells.

Although the current study has revealed an essential role of TRAF4 in ST2^+^ mTh2 cells in the contexts of IL-33-mediated allergen-independent and -dependent type 2 airway inflammation, several questions remain to be addressed. For example, what is the impact of TRAF4 RING-domain E3 activity on IL-33-mediated pathways and functions in ST2^+^ mTh2? It is also interesting to know whether TRAF4 contributes to the steroid-resistant characteristics of ST2^+^ mTh2 given its critical role in promoting cell growth and proliferation. Finally, a comprehensive analysis of IL-33-induced transcriptomes and proteomes altered by TRAF4 deficiency is required to achieve an in-depth understanding of TRAF4’s functions in ST2^+^ mTh2 cells.

## Acknowledgments

This study was supported by the NIH funding (HL144497, HL103453, 5P01CA062220, and 5P01 HL103453)

## Author contributions

C.L. and X.L. conceived and designed all the experiments. J.X., C.L., X.C., W.Q., K.B. L.H., and W.L. performed the experiments and collected the data. C.L. and X.L. wrote the manuscript and supervise the whole project.

---

**Supplemental Figure 1.**
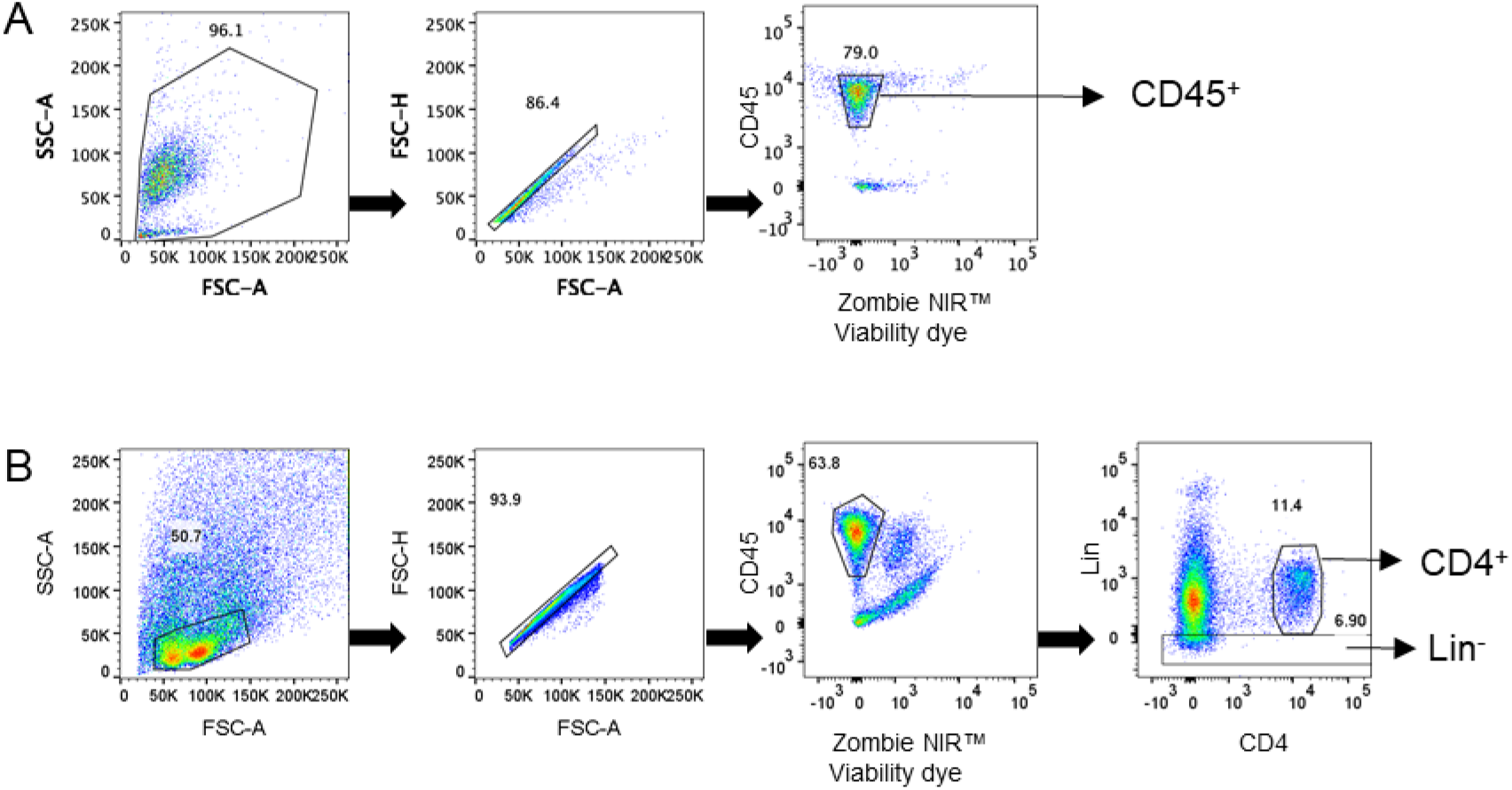
Gating strategies for live CD45^+^ (A), CD4^+^ and lineage negative (Lin^-^) cells (B) in the BAL or lung tissue. BAL cells and single lung cells were first gated by (FSC-A x SSC-A) to remove debris, followed by (FSC-A x FSC-H) to remove doublets, and then the dead cells were excluded using Zombie NIRTM Fixable Viability dye. BAL, Bronchoalveolar lavage.

**Supplemental Figure 2.**
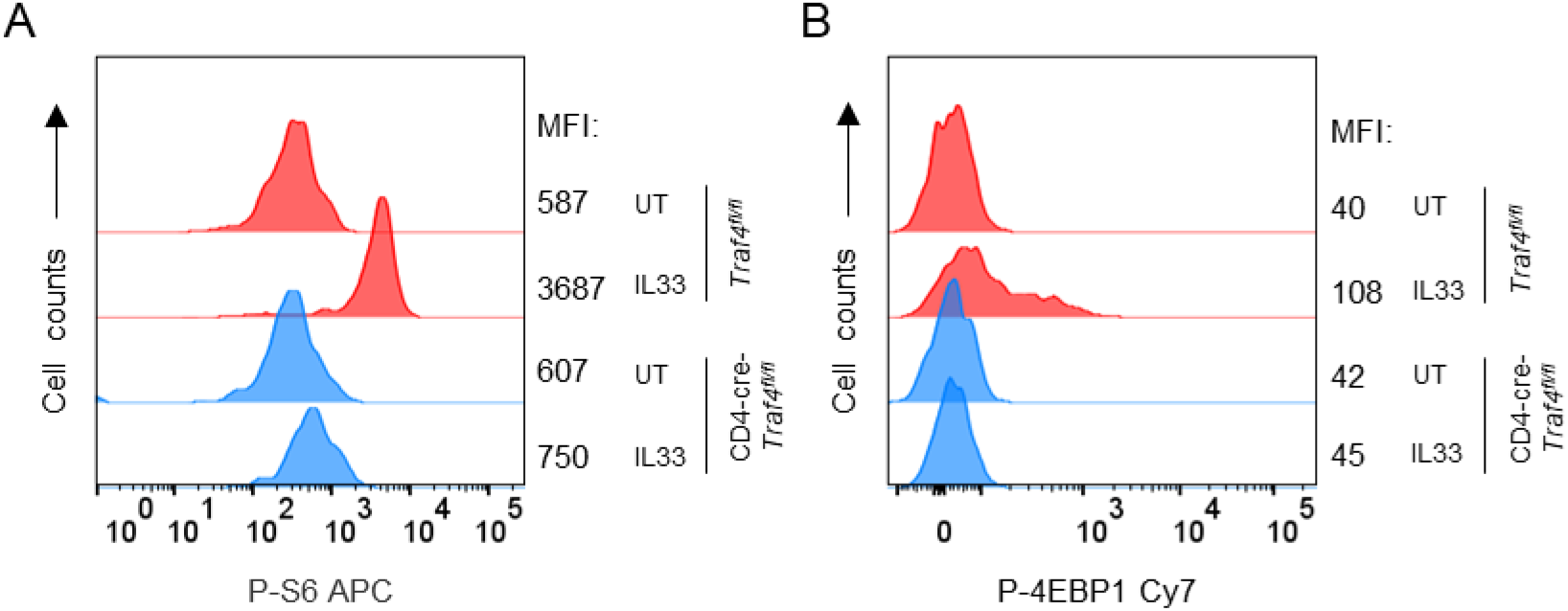
TRAF4 deficiency impairs IL-33-induced phosphorylation of S6 ribosomal protein and 4EBP1. Representative histogram blots showing the surface expression of phospho-S6 ribosomal protein (P-S6) and phospho-4EBP1 (P-4EBP1) on TRAF4-deficient (CD4cve-*Traf4^fl/fl^*) and TRAF4-sufficient (*Traf4^fl/fl^*) mTh2 cells treated with sham or IL-33 for 24 h. Eso, eosinophils. MFI, Mean fluorescence intensity. Plotted data were shown as means ± SEM. All data are representative of two independent experiments.

