## Supplemental figure for "TRAF4 is crucial for the propagation of ST2^+^ memory Th2 cells in IL-33-mediated type 2 airway inflammation"

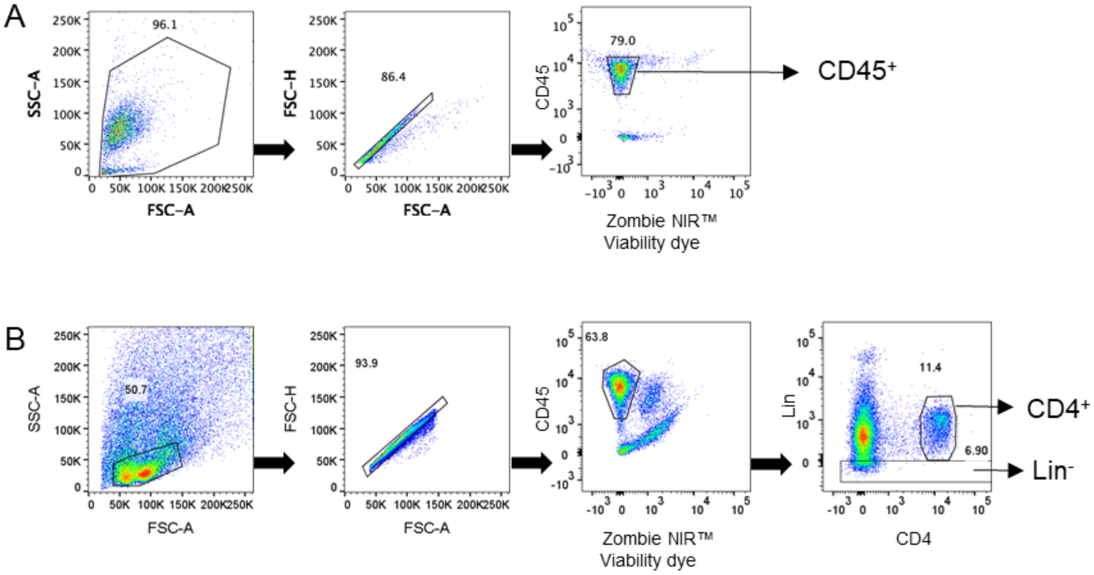


**Supplemental Figure 1**. Gating strategies for live $CD45^+$ (A), $CD4^+$ and lineage negative ($Lin^-$) cells (B) in the BAL or lung tissue. BAL cells and single lung cells were first gated by (FSC-A x SSC-A) to remove debris, followed by (FSC-A x FSC-H) to remove doublets, and then the dead cells were excluded using Zombie NIRTM Fixable Viability dye. BAL, Bronchoalveolar lavage.

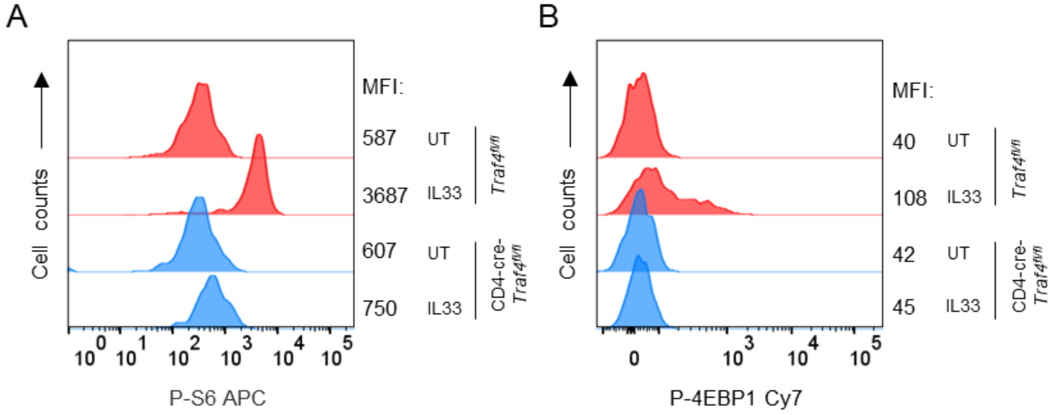


**Supplemental Figure 2.** TRAF4 deficiency impairs IL-33-induced phosphorylation of S6 ribosomal protein and 4EBP1. Representative histogram blots showing the surface expression of phospho-S6 ribosomal protein (P-S6) and phospho-4EBP1 (P-4EBP1) on TRAF4-deficient (CD4cre-*Traf4*$^{fl/fl}$) and TRAF4-sufficient (*Traf4*$^{fl/fl}$) mTh2 cells treated with sham or IL-33 for 24 h. Eso, eosinophils. MFI, Mean fluorescence intensity. Plotted data were shown as means ± SEM. All data are representative of two independent experiments.
